# Effects of bacterial lipopolysaccharide and Shiga Toxin on induced Pluripotent Stem Cell-derived Mesenchymal Stem Cells

**DOI:** 10.1101/2021.09.07.459335

**Authors:** Daiana Martire-Greco, Alejandro La Greca, Luis Castillo Montañez, Celeste Biani, Antonella Lombardi, Federico Birnberg-Weiss, Alessandra Norris, Flavia Sacerdoti, María Marta Amaral, Nahuel Rodrigues-Rodriguez, Jose Ramón Pittaluga, Verónica Alejandra Furmento, Verónica Inés Landoni, Santiago Gabriel Miriuka, Carlos Luzzani, Gabriela Cristina Fernández

**Affiliations:** Laboratorio de Fisiología de los Procesos Inflamatorios. Instituto de Medicina Experimental (IMEX-CONICET). Academia Nacional de Medicina, Buenos Aires, Argentina; Laboratorio de Investigación Aplicada a Neurociencias (LIAN), FLENI-CONICET, Buenos Aires, Argentina; Laboratorio de Fisiopatogenia, Instituto de Fisiología y Biofísica Bernardo Houssay (IFIBIO Houssay-CONICET), Departamento de Fisiología, Facultad de Medicina, Buenos Aires (Argentina); Consejo Nacional de Investigaciones Científicas y Técnicas (CONICET), Buenos Aires, Argentina

**Keywords:** Mesenchymal Stem Cells, Endothelial injury, Shiga-toxin, Lipopolysaccharide, regeneration, Hemolytic Uremic Syndrome

## Abstract

**Background:** Mesenchymal Stem Cells can be activated and respond to different bacterial toxins. Lipopolysaccharides (LPS) and Shiga Toxin (Stx) are the two main bacterial toxins present in Hemolytic Uremic Syndrome (HUS) that cause endothelial damage. In this work we aimed to study the response of iPSC-MSC to LPS and/or Stx and its effect on the restoration of injured endothelial cells.

**Methods:** iPSC-MSC were used as a source of mesenchymal stem cells (MSC) and Human Microvascular Endothelial Cells-1 (HMEC-1) as a source of endothelial cells. iPSC-MSC were treated or not with LPS and/or Stx. For some experiments, Conditioned Media (CM) were collected from each plate and incubated with an anti-Stx antibody to block the direct effect of Stx, or Polymyxin to block the direct effect of LPS. In CM from both treatments, anti-Stx and Polymyxin were used. Results are expressed as mean ± S.E.M. Significant differences (p<0.05) were identified using one way analysis of variance (ANOVA) and Bonferroni’s Multiple comparison test.

**Results:** The results obtained showed that LPS induced a pro-inflammatory profile on iPSC-MSC, but not Stx, even though they expressed Gb_3_ receptor. Moreover, LPS induced on iPSC-MSC an increment in migration and adhesion to gelatin substrate. Also, the addition of CM of iPSC-MSC treated with LPS+Stx, decreased the capacity of HMEC-1 to close a wound, and did not favor the formation of new tubes. Proteomic analysis of iPSC-MSC treated with LPS and/or Stx revealed specific protein secretion patterns that support many of the functional results described here.

**Conclusions:** In conclusion, these results suggest that iPSC-MSC activated by LPS acquired a pro-inflammatory profile that induces migration and adhesion to extracellular matrix proteins (ECM), but the combination LPS+Stx decreased the repair of endothelial damage. The importance of this work is that it provides knowledge to understand the context in which iPSC-MSC could benefit or not the restoration of tissue injury, taking into account that the inflammatory context in response to a particular bacterial toxin is relevant for iPSC-MSC immunomodulation.

## Introduction

Mesenchymal Stem Cells (MSC) are multipotent cells associated with the treatment of different pathologies due to their regenerative properties, thus providing an interesting therapeutic option for various diseases, mainly those that are present in an inflammatory response and tissue damage [1]. They are an heterogeneous subset of stromal stem cells that can be isolated from many adult tissues. However, isolating MSC and obtaining a considerable number for handling often present difficulties. In this sense, derivation of MSC from induced Pluripotent Stem Cells (iPSC) is a widely accepted method that results in cells that have similar properties to those obtained directly from adult tissues. In this sense, our group described in a previous paper a robust and fast method to obtain the iPSC-MSC that were used in this work [2].

In recent years, the use of MSC has increased in important clinical applications [1]. It has been described that these cells are involved in immune processes and participate in the repair of many types of tissue injuries, mainly in a paracrine fashion by secreting numerous soluble factors [3, 4]. Also, it is widely reported the capacity of MSC to migrate into injured sites and to release inflammatory and growth factors [5]. Some types of MSC, like those obtained from bone marrow, can potentially move from their niche into the circulation crossing through endothelial cells from vessel walls to the site of damage and adhere to the extracellular matrix (ECM) around the wound. However, the trafficking of MSC from their niche to target tissues is a complex process. The migration process is affected by chemokines, cytokines, growth factors and mechanical factors such as shear stress, vascular cyclic stretching, and ECM adhesion [6].

Another important aspect of MSC is their influence on different functions of surrounding cells in order to repair tissue damage by promoting migration, adhesion and differentiation, also determining cellular and biochemical changes in all phases of tissue damage [1]. MSC also plays a role in immune processes as they can respond to bacterial toxins and inflammatory cytokines. In this sense, it has been described that MSC can be polarized in vitro towards either pro- or anti-inflammatory phenotypes, depending on the Toll-Like Receptor (TLR) ligand involved in their activation [7, 8]. In many infections TLR4 are activated with lipopolysaccharides (LPS) which are endotoxins present on the outer surface of Gram Negative bacteria, and stimulate MSC toward a pro-inflammatory phenotype, modulating some of their functions [10]. Also, these danger signals activate immune cells, and start an appropriate host response with the aim to reestablish homeostasis by recruiting them to the site of injury. However, if the inflammatory response turns out to be excessive, tissue damage repairment may not be possible [1]. Furthermore, TLRs are crucial in sensing signals and switching immune responses from MSC depending on the inflammatory state, contributing in this way with the immunomodulatory properties that MSC are well known to have [9].

Hemolytic uremic syndrome (HUS) is a disease caused by infections with enterohemorrhagic Gram-negative bacteria that produce Shiga toxin (Stx). Once Stx accesses the circulation, it interacts with a globotriaosylceramide glycolipid receptor (Gb_3_) in target cells and this interaction leads to a cascade of events that usually culminates with the inhibition of protein synthesis and cell death. Endothelial cell damage is a central event in the pathophysiology of HUS and is the most important factor of the microangiopathic process typically found in this disease [12]. In particular, glomerular endothelial injury triggers a thrombotic microangiopathy leading to the formation of platelet and fibrin thrombi that occlude the microvasculature of the glomerulus, affecting renal function and resulting in acute renal failure. In addition to the toxic effects caused by the interaction of Stx with target cells, *in vivo* and *in vitro* evidence have demonstrated that LPS, present in all Gram-negative bacterial infections, potentiate endothelial cell damage by increasing susceptibility of these cells to the toxin [13, 14, 15]. Moreover, the presence of LPS triggers a strong inflammatory response, which can also contribute to endothelial dysfunction [16].

Taking into account that in many infections iPSC-MSC can be activated due to bacterial toxins and this can be decisive to reestablish homeostasis, the aim of this work was to investigate whether LPS and/or Stx treatments modify the iPSC-MSC secretory profile, and/or functions that could modulate the characteristic endothelial damage induced in the context of HUS.

## Materials and Methods

### Cell cultures and treatments

iPSC-MSC were obtained and differentiated as previously published [2]. These cells were maintained using alpha-MEM medium (Gibco, Ireland) supplemented with platelet lysate, 10 % of Penicillin-Streptomycin and glutamine (Gibco, Ireland). At 80 % of confluence, cells were trypsinized with 0.25 % of trypsin-EDTA (Gibco, Ireland). Human dermal Microvascular Endothelial Cells-1 (HMEC-1) were used to perform the experiments of endothelial damage. Cells were cultured at 37°C in a 5 % CO_2_ humidified atmosphere using MCDB-131 medium (Gibco, Ireland) with phenol red and supplemented with 15 % fetal bovine serum (Natocor, Argentina), penicillin (Gibco, Ireland), streptomycin (Gibco, Ireland), L-glutamine 10 mM (Gibco,Ireland), hydrocortisone 1 *μ*g/mL (Sigma, USA) and endothelial cell growth supplement 20 *μ*g/mL (Abcys, France). We set 4 treatment groups: Control (vehicle: only alpha-MEM), LPS (Sigma, USA, 0,5 ng/ml), Stx (Toxin Technology, USA, 20 ng/ml) and LPS+Stx. We exposed iPSC-MSCs and endothelial cells to Stx for 24 h. We used the type 2 variant of Stx (Stx2), as it is the most relevant in terms of epidemiology [17]. Also, as LPS is present in all Gram negative bacterial infections, and is the principal modulator of the inflammatory response, we use the combination of LPS+Stx for every experiment. Both LPS or Stx were added at the same time in LPS+Stx treatments. Also, Polymyxin B (Sigma, USA) was used in all treatments with Stx in order to avoid LPS contamination.

### Conditioned Media

iPSC-MSC were seeded in alpha-MEM and treated with LPS and/or Stx during 24 h. Then, conditioned media (CM) were collected and incubated for 2 h with an anti-Stx antibody (anti-Stx2 variant from Toxin Technology, USA) to block the direct effect of Stx, or Polymyxin (Sigma, USA) to block the direct effect of LPS. In LPS+Stx CM, both anti-Stx and Polymyxin were used.

### Viability assays

Cells were plated at subconfluency, treated for 24 h, and then gently washed to remove dead cells. After that, the remaining attached cells were fixed and dyed for 20 min using a solution of 0.1 % crystal violet in 20 % methanol. Then, the crystals were solubilized with 30 % acetic acid and measured with an ELISA detector at 540 nm. Several washes were done in order to eliminate the residual dye.

### Proliferation

1 x 10^5^ iPSC-MSC cells were seeded in 96 well plates with LPS and/or Stx for 48 h at 37°C in 5 % CO_2_. Then 0.5 *μ*Ci/well of 3H-thymidine was added and incubated for another 20 h. After that, cells were harvested, scintillation fluid was added, and the radioactive thymidine incorporated into DNA was measured.

### Migration / Scratch assays

iPSC-MSC or HMEC-1 cells were seeded to confluence in 24-well plates (Jet Biofil) in the corresponding culture media. In the case of iPSC-MSC (migration assays), cells were incubated for 24 h with media containing either Control or toxins (LPS, Stx, LPS+Stx) before scratch, while in HMEC-1 cells (wound repair) the conditioned media from treated iPSC-MSC was added immediately after doing the scratch. Starting point (time 0) of the experiment was defined as the moment when cells were returned to the incubator, with an end point of 18 h. Images were captured at both instances with a Nikon Eclipse T5 100 microscope and then analyzed with ImageJ software. We used the freehand tool to manually draw over the gap edges to determine the area of the wound at time 0, 6 h (for iPSC-MSC) or 18 h (for HMEC-1). The percentage of gap closure was calculated as: [(gap area at 0 h - gap area at x h)/gap area at 0 h] x100.

### Adhesion assay

Adhesion of iPSC-MSC was evaluated on 96-well plates previously coated with 2 % gelatin (40 min at room temperature, Sigma, USA). Cells were first treated with vehicle (Control), LPS, Stx and LPS+Stx and then collected with trypsin, counted and seeded in the gelatin-coated wells (20,000 cells/well). Cells were allowed to attach for 15 min at 37°C and stained with crystal violet solution. Images of adhered cells were captured using a Nikon Eclipse T5 100 microscope and quantified with the ImageJ software using the count cell option.

### Gb_3_ measurement by Thin Layer Chromatography (TLC)

Gb_3_ levels were detected by TLC and analyzed by densitometry. iPSC-MSC cells were cultured in flasks and grown at 37°C in an atmosphere of 5 % CO_2_ until cells were nearly confluent. Cells were treated with Stx and/or LPS as previously described. From each treatment, total cells glycolipids were extracted according to the method of Bligh and Dyer et. al [18]. Briefly, 3 ml of chloroform:methanol 2:1 v/v were incorporated into the cells, and during 15 min cells were incubated on ice. Two ml of chloroform:water (1:1) were added and centrifuged at 3,000 rpm for 5 min to separate phases. The lower phase, corresponding to the neutral glycolipid extract, was brought to dryness and used for Gb_3_ determination. One ml of methanol and 0.1 ml of 1.0 M NaOH was added to the dried residue, and incubated 16 h at 37°C. Fractionated lipids were subjected to TLC with a silica gel 60 aluminum plate previously activated by incubation 15 min at 100°C, in a glass tank with a mixture of chloroform, methanol, and water (65:35:8). To compare quantities, a purified glycosphingolipid standard (0.5-1 and 2 *μ*g) (Matreya, USA) was also added to the plate. After the solvent front reached the top of the plate, the gel matrix was air dried and treated with a solution of orcinol, water and sulfuric acid (Acros Organics, USA) to visualize the separated carbohydrate and glycolipid components. The densitometric analysis of Gb_3_ bands was analyzed by Image Quant 5.0 software. Values are expressed as ng of Gb_3_/10^6^ cells.

### ELISA assays

Detection of TNF-*α* (BioLegend cat. 430205), IL-8 (BioLegend cat. 78141), TGF-*β* (Biolegend, cat. 436707), and IL-10 (Biolegend, cat. 430601) from iPSC-MSC were performed with ELISA kits, following manufacturer’s recommendations. Concentration results were obtained in pg/ml.

### Tubulogenesis assay

Assays were performed on 96-well plates coated with geltrex at 37°C for no less than 30 min. Approximately 15,000 HMEC-1 cells/100 *μ*l were seeded on coated wells using EGM-2 media (Lonza, Switzerland) and incubated overnight either with conditioned media from toxin-treated iPSC-MSC or vehicle at 37 °C. Images of tube formation were captured the following day (24 h) using a Nikon Eclipse T5 100 microscope followed by analysis on ImageJ software using the count the branch points option obtained.

### Mass spectrometry (MS)

LC-MS/MS (Liquid Chromatography with tandem mass spectrometry) assays and MS analysis on Conditioned Media of three technical replicates from vehicle (Control), LPS, LPS+Stx and Stx-treated iPSC-MSC were performed at the Proteomics Core Facility CEQUIBIEM (University of Buenos Aires, Buenos Aires, Argentina) following specifications detailed in La Greca et al, 2018 [19]. Briefly, peptides were reduced with dithiothreitol (DTT), precipitated with trichloroacetic acid (TCA) and digested with trypsin. Approximately, 1 *μ*g of protein digests were analyzed by nanoLC-MS/MS in a Thermo Scientific QExactive Mass Spectrometer coupled to a nanoHPLC EASY-nLC 1000. Data acquisition and configuration for peptide identification were achieved with XCalibur 3.0.63 software and raw data produced was fed into Proteome Discoverer software to classify identified peptides against Homo sapiens protein sequences database (trypsin specificity) and quantify abundance (area under the curve strategy).

### Bioinformatic analysis of MS data

Protein abundance obtained from Proteome Discoverer in the form of area-based quantification (area under the curve) [20] was employed for downstream bioinformatic analysis. Technical replicates were collapsed and samples normalized by total area (total area per sample/1000) using custom Python scripting. Peptide abundance identified as ALBUMIN (P02768) was excluded from further analysis as it is most likely a residual contaminant from the platelet lysate used during iPSC-MSC routine culture. The rest of the identified proteins were clustered using a hierarchical-based approach and plotted in a heatmap using pheatmap package in R. In order to aid visualization of identified proteins, ProteinIDs were mapped to Gene Names using the uniprotID converter (www.uniprot.org). Ontological terms classified as “Biological processes” (BPs) were determined using DOSE [21] and clusterProfiler [22] packages keeping only the top ten statistically significant over-represented terms (p-value<0.01, q-value<0.05). These over-represented BPs - also called enriched-were determined by statistically testing (Fisher’s exact test followed by hypergeometric distribution test to evalu-ate significance) the relationship between the frequency of genes/proteins present in any given term (observed or sample frequency) and the frequency of genes/proteins annotated to the same term (expected or background frequency). Ultimately, this means that enriched BPs showed observed frequency values higher than their expected frequency for that term, and the difference proved to be significant (p-value<0.01).

### Statistical analysis

Results are expressed as mean ± S.E.M. Significant differences (p<0,05) were identified using one way analysis of variance (ANOVA) and Bonferroni’s Multiple Test Comparison using GraphPad software package (Prism 5.0 Version, San Diego, USA).

## Results

### Gb_3_-expressing iPSC-MSCs remained viable after LPS and Stx treatments

In order to establish the concentrations of Stx and LPS to be used with iPSC-MSC and endothelial cells we set two dose response curves with different concentrations. As a first step in determining the Stx concentration needed to cause endothelial damage, we treated Human Microvascular Endothelial Cells (HMEC-1) with different doses of this toxin, and measured the resultant viability after 24 h. As shown in Figure 1A, we found that concentrations of 5, 10 and 20 ng/ml of Stx were sufficient to cause endothelial cell death in a dose dependent manner alone. LPS did not show any additional toxic effect when combined with Stx. Although LPS alone did not induce HMEC-1 cytotoxicity, this concentration of LPS was able to modulate endothelial cell functions, by increasing ICAM-1 expression (Supp. Figure S1). In contrast to the results observed in endothelial cells, none of the concentrations of Stx or Stx in combination with LPS (LPS+Stx) affected iPSC-MSC viability (Figure 1B), or their proliferation measured by 3H-thymidine incorporation (Figure 1C).

**Figure 1:**
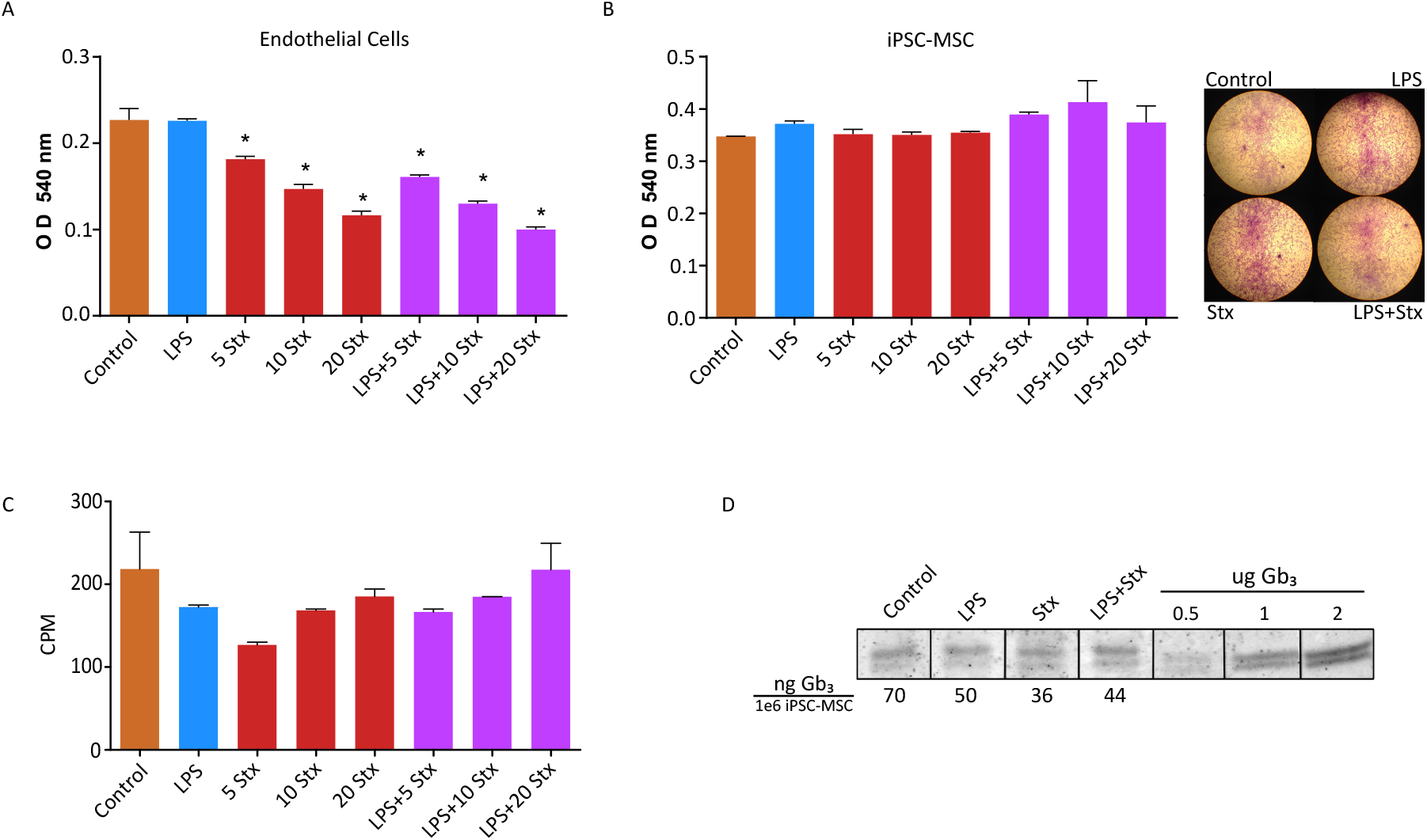
Effect of Stx and LPS on the viability of endothelial cells, iPSC-MSC and its Gb_3_ expression. (A) Viability of HMEC-1 treated with LPS (500 ng/ml), alone or in combination with different Stx concentrations (5-20 ng/ml) were measured at 540 nm. and optic density (O.D.) from crystal violet were represented from each treatment. (B) Viability of iPSC-MSC treated with LPS (500 ng/ml), alone or in combination with different Stx concentrations (5-20 ng/ml) were measured at 540 nm. and optic density (O.D.) from crystal violet were represented from each treatment. Representative microphotographs depicting iPSC-MSC cultures are shown in the right panel (x10). (C) Proliferation was measured by 3H-thymidine incorporation on control and treated iPSC-MSC, 72 h post stimulus. Counts per minute (CMP) are shown. (D) Thin Layer Chromatography (TLC) assays performed in Control and treated iPSC-MSC to measure the expression of Gb3 receptor compared to known Gb_3_ standards (0.5, 1 and 2 *μ*g). Gb_3_ quantification (ng Gb_3_/1e^6^ iPSC-MSC) is shown below each treatment column. Results were expressed as mean ± S.E.M. n = 12–18 per group; *P<0.05.

In addition, because Stx did not affect iPSC-MSC viability or proliferation levels, we decided to determine the presence of the Gb_3_ receptor in these cells. We obtained similar levels of Gb_3_ expression on iPSC-MSC in both Control and treated conditions (LPS, Stx, and LPS+Stx) using thin layer chromatography (Figure 1D).

These results indicate that even though iPSC-MSC expresses the Stx receptor, treatment with this toxin alone or in combination with LPS does not affect cell viability, in contrast to what was observed for endothelial cells.

### LPS induced a pro-inflammatory program on iPSC-MSC but not Stx

iPSC-MSC regulate their microenvironment releasing different cytokines that can modulate biological processes in an autocrine or paracrine way [23]. Moreover, inflammatory signals released in many infections are associated with migration, adhesion to the extracellular matrix and many mechanisms near the site of inflammation [10].Therefore, in order to determine the immunomodulatory contribution of LPS- or Stx-treated iPSC-MSC, we measured the release of the pro-inflammatory cytokines IL-8, and TNF-*α*, and the antiinflammatory cytokines TGF-*β* and Il-10. As shown in Figure 2A and B, only LPS significantly increased the release of Il-8 and TNF-*α* compared to Control cells. The addition of Stx alone did not induce the release of IL-8 or TNF-*α*. Also, when iPSC-MSC were exposed to LPS+Stx, they increased the production of Il-8 and TNF-*α* compared to basal cells, but TNF-*α* levels were lower compared to LPS alone. Additionally, the presence of LPS, Stx or LPS+Stx decreased significantly the levels of the TGF-*β* compared to control cells. Moreover, both Il-10 and VEGF levels remained undetectable in all conditions (<15.6 and <31.3 pg/ml which are the lower detectable concentrations with ElISA kits respectively). These results indicate that LPS polarizes iPSC-MSC towards a pro-inflammatory phenotype, but Stx does not contribute to this polarization despite being a bacterial toxin.

**Figure 2:**
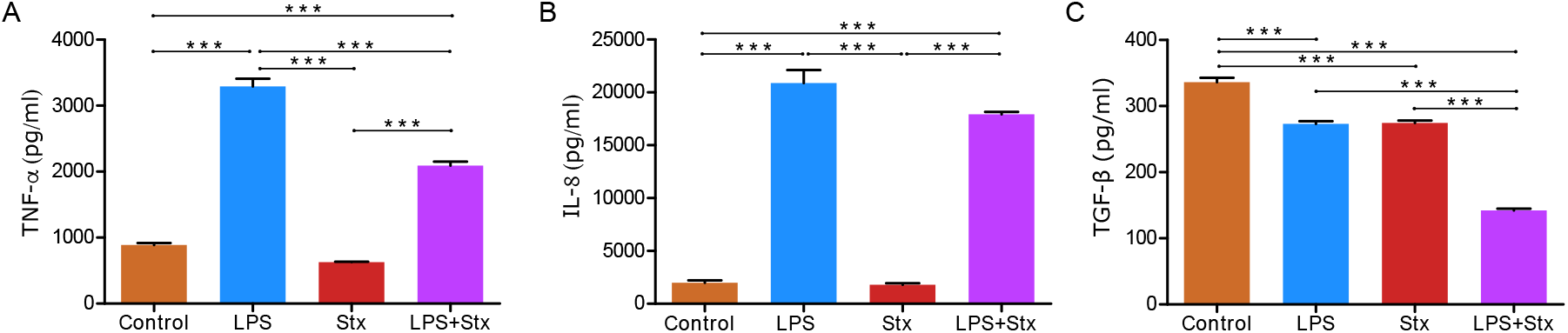
Inflammatory cytokines are produced by iPSC-MSC in contact with LPS. Stx did not have this effect. iPSC-MSC were treated with LPS and/or Stx for 24 h and then secreted TNF-*α* (A), IL-8 (B) and TGF-*β* (C) were determined using ELISA kits. Results were expressed as mean ± S.E.M. n = 3 per group; ***P<0.001.

### LPS and not Stx increased the migration of iPSC-MSC

It is known that in some inflammatory pathologies, iPSC-MSC can respond to a wide range of extracellular signals and modulate some of their functions [10, 9]. In this sense, we investigated the effect of LPS and Stx on the capacity of iPSC-MSC to migrate after a wound was performed. First, cells were incubated with LPS, Stx or a combination of both toxins for 24 h, and then a scratch was performed mechanically across the cell monolayer. We observed that LPS treatment increased the percentage of migration of iPSC-MSC compared to control cells (Figure 3). Conversely, Stx did not modify this function showing similar migration as in basal condition. The combination of LPS+Stx increases these percentages similar to LPS alone when compared to control and Stx treated cells. In conclusion, the inflammatory stimulus LPS increases the migration of iPSC-MSC, whereas Stx does not modify this effect.

**Figure 3:**
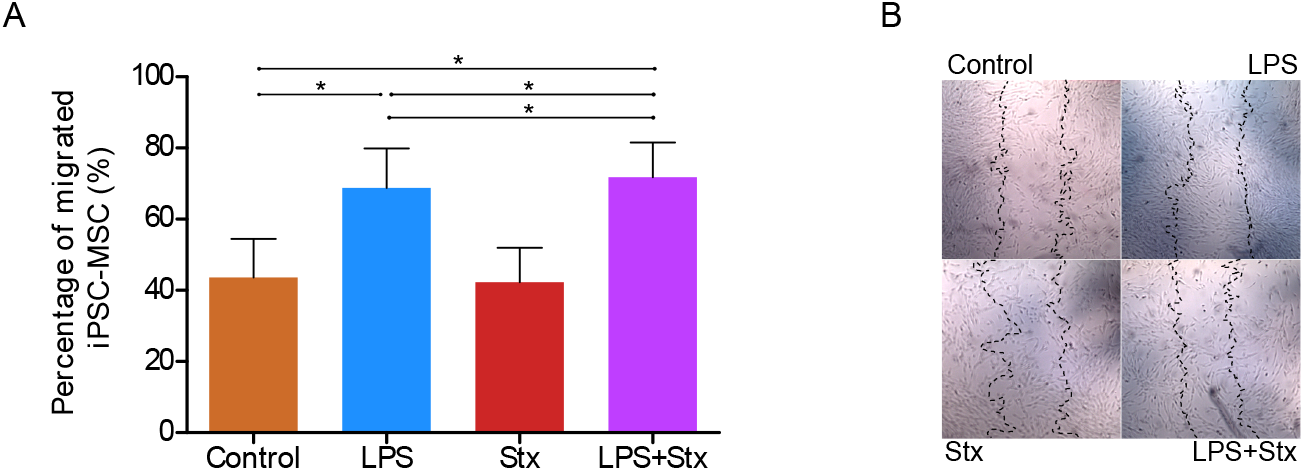
LPS increased iPSC-MSC migration and Stx did not modulate this function. Percentage of migrated area from LPS and/or Stx treated or Control iPSC-MSC after 6 h post-scratch over the monolayer of the cells.Representative microphotographs are shown for each treatment in the right panel. The discontinuous line represents the wound at time 0. Results were expressed as mean ± S.E.M. n = 8 per group; *P<0.05.

### The combination of LPS+Stx augmented iPSC-MSC capacity to adhere to extracellular matrix

The fact that iPSC-MSC migrate sensing inflammatory signals involves an adhesion to the extracellular matrix (ECM) in order to reach the site of damage [24]. In this sense, iPSC-MSC were used for measuring adhesion to a substrate (gelatin), 24 h after incubation with LPS, Stx or LPS+Stx. Treated cells were harvested and settled on gelatine covered wells. After 15 minutes non-adhered cells were eliminated by vigorous washing and remnant gelatin-adhered cells were stained with crystal violet and counted by microscopy. We found that the treatment with LPS or Stx alone did not alter adhesion to gelatin, although a slight increase on adherent cells was found. The combination of both toxins (LPS+Stx) caused a statistically significant increase in cell adhesion (Figure 4). This result suggests that the effect of LPS on iPSC-MSC adhesion to a gelatin matrix is potentiated by the presence of Stx.

**Figure 4:**
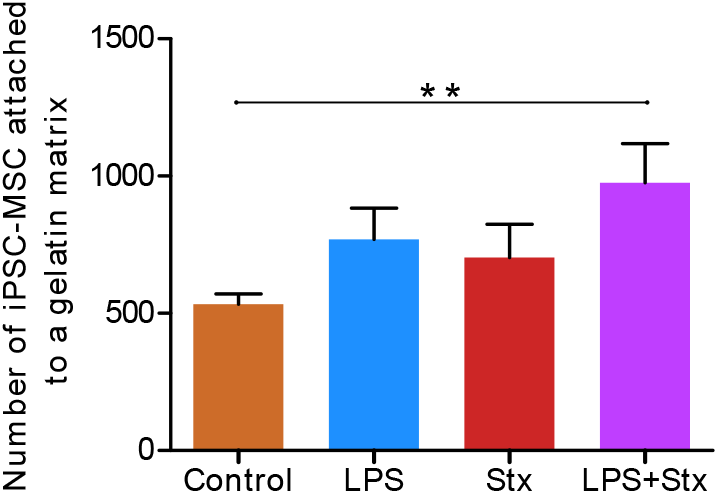
LPS+Stx increased in iPSC-MSC the adhesion to gelatin. Adhered iPSC-MSC to gelatin after 24 h of being treated with LPS and/or Stx. Results were expressed as mean ± S.E.M. n = 4 per group; **P<0.01.

### Conditioned media (CM) from iPSC-MSC exposed to LPS+Stx decreased the capacity to repair endothelial damage

In order to investigate the effect of pro-inflammatory iPSC-MSC on endothelial repair, we performed a wound healing assay. For this purpose, a scratch was performed across a monolayer of endothelial cells HMEC-1. Then, cells were treated with CM obtained from iPSC-MSC that have been previously treated or not with LPS, Stx, LPS+Stx for 24 h. Then, the percentage of endothelial wound repair was measured. Figure 5A shows that the presence of CM from LPS+Stx-treated iPSC-MSC reduced wound closure compared to non-treated iPSC-MSC CM. CM from LPS and Stx alone-treated iPSC-MSC did not affect this function. Moreover, none of the CM affected the formation of new tubes on endothelial cells (Figure 5B). These results indicate that the effect of Stx and LPS treatment on iPSC-MSC does not favor the repair of endothelial damage.

**Figure 5:**
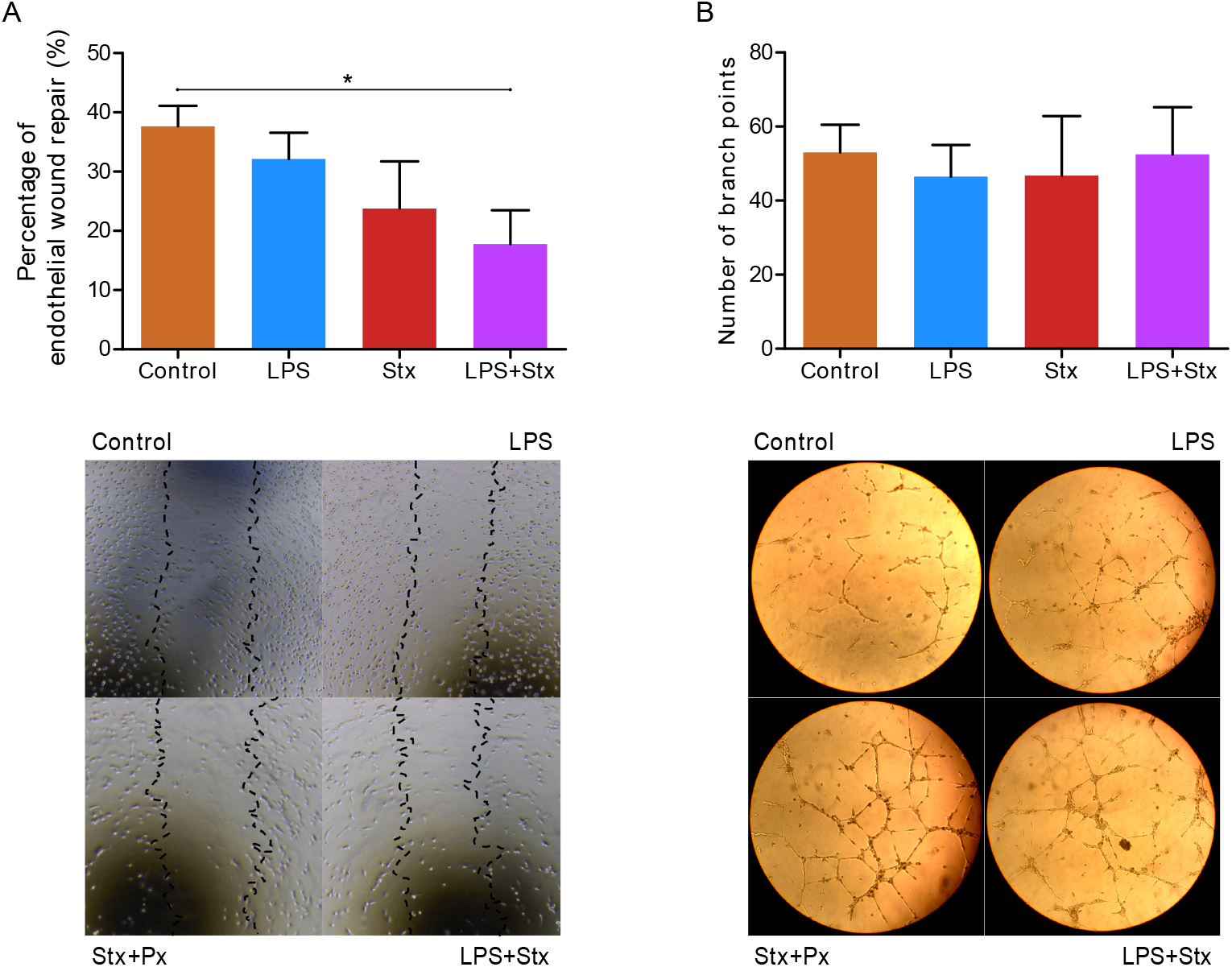
LPS+Stx decreased in iPSC-MSC repair mechanisms in endothelial cells. Conditioned media (CM) from iPSC-MSC treated with LPS and/or Stx were added to endothelial cells for (A) wound healing assay (percentage of endothelial wound repair is shown and representatives microphotographs are shown) (B) tubulogenesis assay (number of branch points is shown and representatives microphotographs are shown). Results were expressed as mean ± S.E.M. n = 8 per group; *P<0.05.

### Analysis of proteins secreted by iPSC-MSC treated with LPS and or Stx

With the objective to explore the proteins secreted by iPSC-MSC in the CM after the treatments with LPS and/or Stx, we performed a proteomic analysis, as this technique allows for the simultaneous identification of the proteins present in any given sample, providing a useful and fast method to assess relevant pathways or biological processes [25]. Thus, we studied the expression levels of the proteins secreted to the CM by untreated cells (Control) or cells treated with LPS, Stx and LPS+Stx iPSC-MSC. Hierarchical clustering of protein abundance data produced four different groups revealing specific secretion profiles associated with each experimental condition (Supp. Figure S2A). Functional analysis on clustered data resulted in a set of over-represented ontological terms (Supp. Figure S2B), in the form of biological processes (BPs), that exposed important aspects of bacterial toxin treatment.

Gene ontology over representation analysis on clustered data showed that some proteins are more represented in the CM from iPSC-MSC after treatment with LPS when compared to control cells or with Stx and LPS+Stx treatments. The proteins found in the CM from iPSC-MSC treated with LPS but not with the combination of both toxins are related to BPs like “acute inflammatory response”, “platelet degranulation”, “regulation of fibrinolysis” and “extracellular matrix organization” (e.g. SERPINE1, AHSG1, FN, THBS1, PLG, PTX3 and CCN2 (Figura 6A)), in line with results obtained in Figure 2 where LPS polarized iPSC-MSC to a pro-inflammatory profile increasing migration and adhesion to extracellular matrix (Figure 3 and 4).

Furthermore, LPS+Stx increased the expression of proteins related to BP “IL-12 mediated signaling pathway”, “endopeptidase activity” and “actin filament organization” (e.g. PPIA) (Figure 6B). These proteins can be associated with Figure 5, where a decreased capacity of wound closure was observed in endothelial cells incubated with the iPSC-MS treated with LPS+Stx.

**Figure 6:**
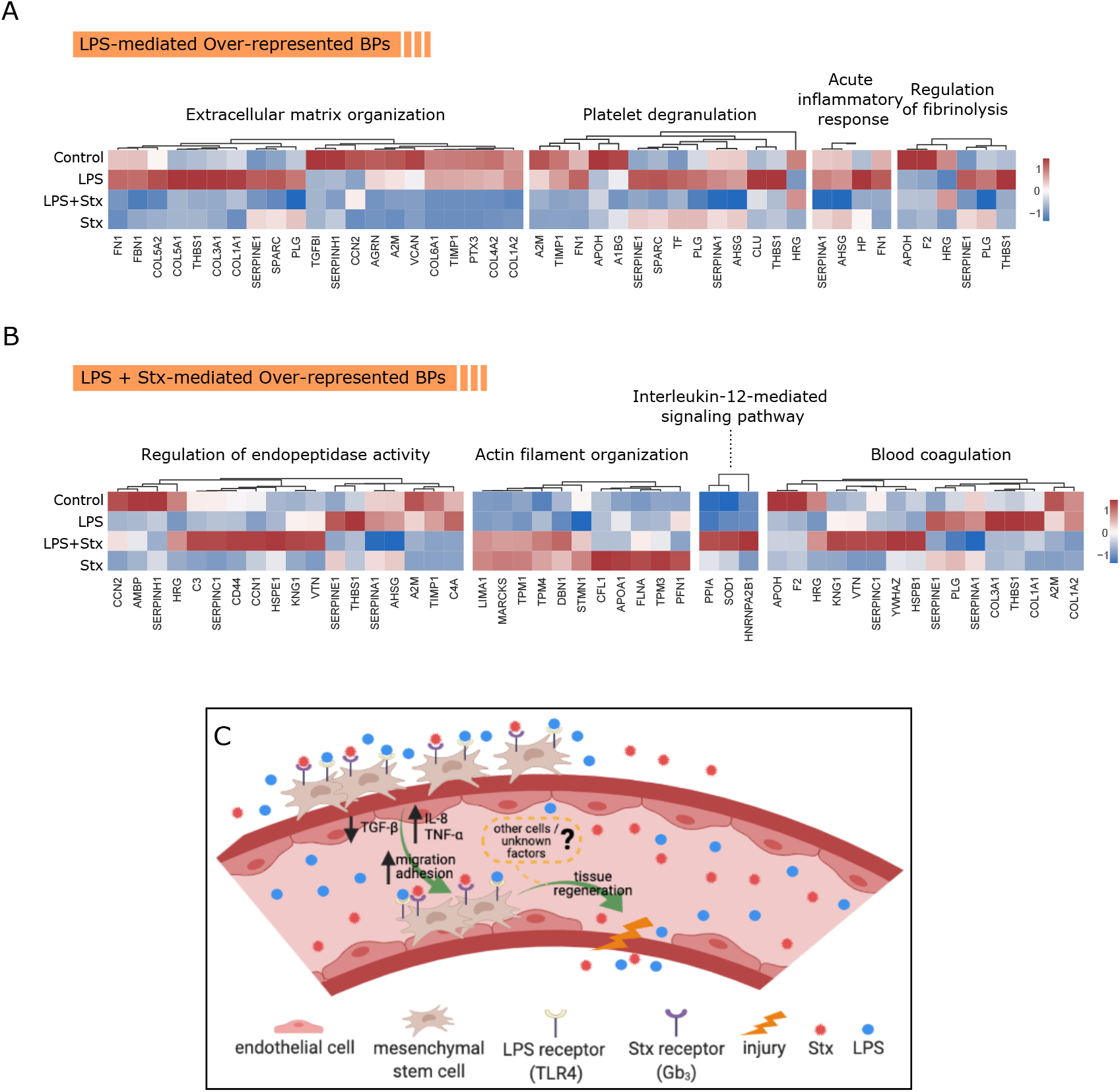
Protein abundance levels found in over-represented biological processes from toxin-treated iPSC-MSC. Comparative protein levels among treatments for biological processes found predominantly in (A) clusters 3 and 4 and in (B) clusters 1 and 2. Heatmap was plotted using scaled (z-score) normalized areas in which red color indicates higher abundance while blue represents low abundance. Dendrograms on top of heatmaps reflect hierarchical clustering of proteins. (C) Schematic representation of a segment of a blood vessel and the events triggered by LPS and Stx treatments focusing on MSC response. Created with BioRender (biorender.com).

## Discussion

Immune responses against bacterial toxins are crucial to resolve infectious pathologies. Usually, different cell types are recruited to respond and participate in order to reestablish the altered homeostasis. The mechanisms involved in these processes include the secretion of a wide range of cytokines to the environment in order to attract different cells that can positively or negatively modulate tissue damage. Mesenchymal stem cells (MSC) are known to participate in these processes secreting cytokines that are involved in many mechanisms with the aim of restoring tissue injury, often present due to infections [10]. In this sense, host immune cells can recognize some bacterial toxins such as LPS and mount defenses to clear pathogens, releasing pro-inflammatory cytokines that contribute to activate immune and non immune cells with the objective to restore homeostasis. TLR4 recognizes LPS, and cell activation through this receptor leads to profound cellular and systemic responses that mobilize innate and adaptive host immune cells [26]. Another bacterial toxin that participates in inflammatory processes is Shiga toxin (Stx). This multifunctional toxin is capable of inducing cell stress and activating innate immune responses that may lead to inflammation increasing the severity of organ injury in HUS patients [27]. Taking this into account, we investigated the effect of LPS and/or Stx on iPSC-MSC. Particularly, in this work it was studied the induction of secreted soluble factors from iPSC-MSC exposed to both bacterial toxins and their possible contribution on endothelial damage.

To the best of our knowledge, this is the first report describing expression of Gb_3_ in iPSC-MSC. This result is relevant for featuring iPSC-MSC as direct potential cellular targets for Stx. It has been demonstrated that Gb3 mediates the entrance of Stx to target cells, e.g. endothelial cells, generally causing cell death [28]. However, we did not observe any toxic effect on iPSC-MSC after incubation with Stx, in contrast to what happens with endothelial cells. In line with this result, Geelen et. al. showed that although monocytes express a receptor for Stx, they do not show any cytotoxic effect after incubation with Stx [29], indicating that cell death is not the only possible result after Stx interacts with its receptor. Furthermore, we observed that iPSC-MSC cultured with LPS increased their capacity to migrate and adhere to extracellular matrix (ECM) proteins. Interestingly, when we measured the ability of iPSC-MSC to attach to the ECM, we observed an additive effect between LPS and Stx in this function, probably reflecting the physiopathological events that occur in a context of infection with a Stx-producing *E. coli* in HUS. These functional results (Figure 3 and 4) were consistent with the secreted proteins obtained in the proteomic analysis from the CM of iPSC-MSC treated with LPS. The results showed that iPSC-MSC contributed to creating an inflammatory environment as we observed in ELISA assays (IL-8 and TNF-*α* increments, Figure 2) but also in the proteomic analysis. For example SER-PINE1 is involved in acute inflammatory responses as is described in hepatocytes, monocytes, macrophages and bronchiolar cells [30], AHSG1 is a protein associated with inflammation and chronic diseases such as endotoxemia and sepsis [31], THBS1 represents a potent pro-inflammatory signal for macrophages, and is also produced by them [32], PLG is an enzyme with a crucial role in inflammation and coagulation [33], FN is a ubiquitous and essential component of the extracellular matrix that participates in many events related to cell migration and adhesion [34] and PTX3 is a prototypic soluble pattern recognition receptor, expressed at sites of inflammation and involved in regulation of tissue homeostasis. Systemic levels of PTX3 increase in many (but not all) immune-mediated inflammatory conditions [35]. All these proteins were found in higher quantities in the CM of iPSC-MSC treated with LPS, in accordance with an induction of the pro-inflammatory program that these cells acquired.

To the best of our knowledge, this is the first report describing the expression of Gb_3_ in iPSC-MSC. This result is relevant for featuring iPSC-MSC as direct potential cellular targets for Stx. It has been demonstrated that Gb3 mediates the entrance of Stx to target cells, e.g. endothelial cells, generally causing cell death [28]. However, we did not observe any toxic effect on iPSC-MSC after incubation with Stx, in contrast to what happens with endothelial cells. In line with this result, Monnens et. al. [29] showed that although monocytes express a receptor for Stx, they do not show any cytotoxic effect after incubation with Stx, indicating that cell death is not the only possible result after Stx interacts with its receptor. Furthermore, we observed that iPSC-MSC stimulated with LPS increased their capacity to migrate and adhere to extracellular matrix (ECM) proteins. Interestingly, when we measured the ability of iPSC-MSC to attach to the ECM, we observed an additive effect between LPS and Stx in this function, probably reflecting the physiopathological events that occur in a context of infection with a Stx-producing *E. coli* in HUS. These functional results (Figure 3 and 4) were consistent with the secreted proteins obtained in the proteomic analysis from the CM of iPSC-MSC treated with LPS. The results showed that iPSC-MSC contributed to creating an inflammatory environment as we observed in ELISA assays (IL-8 and TNF-*α* increments, Figure 2) but also in the proteomic analysis. For example SER-PINE1 is involved in acute inflammatory responses as is described in hepatocytes, monocytes, macrophages and bronchiolar cells [30], AHSG1 is a protein associated with inflammation and chronic diseases such as endotoxemia and sepsis [31], THBS1 represents a potent pro-inflammatory signal for macrophages, and is also produced by them [32], PLG is an enzyme with a crucial role in inflammation and coagulation [33], FN is a ubiquitous and essential component of the extracellular matrix that participates in many events related to cell migration and adhesion [34] and PTX3 is a prototypic soluble pattern recognition receptor, expressed at sites of inflammation and involved in regulation of tissue homeostasis. Systemic levels of PTX3 increase in many (but not all) immune-mediated inflammatory conditions [35]. All these proteins were found in higher quantities in the CM of iPSC-MSC treated with LPS, in accordance with an induction of the pro-inflammatory program that these cells acquired.

Another fact observed in this work was that Stx treatment did not modulate any of the iPSC-MSC functions assayed, and did not generate a pro-inflammatory profile as LPS did, indicating that LPS is the main inducer of a pro-inflammatory profile in these cells in a context of HUS.

Although we did not observe an increase in the percentage of wound repair in endothelial cells exposed to CM from iPSC-MSC treated with LPS+Stx, the combination of both toxins decreased the capacity of iPSC-MSC to restore the endothelial damage and also, did not modify the mechanism of new tube formation. We hypothesize that the treatments with LPS+Stx on iPSC-MSC induce the release of some factors that decrease the capacity of endothelial cells to repair a wound. In this sense, in the proteomic analysis we found that the use of LPS+Stx in iPSC-MSC, induced the release of proteins related to the BP of “IL-12 mediated signalling pathway” such as PPIA, which is reported to promote apoptosis in endothelial cells and chemotaxis in inflammatory cells [36]. As Wong and Waterman et.al described in a previous work, we can speculate that activation of TLR4 in iPSC-MSC due to LPS resulted in the secretion of pro-inflammatory factors in the CM that are important for early injury responses, like migration and cell adhesion to ECM, but probably not to resolve tissue damage [1, 10]. However, these mechanisms could prepare the microenvironment for later iPSC-MSC anti-inflammatory responses that could facilitate the restoration of tissue injury. This second response could be possible as it is known that MSC can promote or inhibit immune responses due to their immunomodulatory properties, determined by the strength of the inflammatory milieu ([9], Figure 6C). In this sense, proteomics results in concordance with the Biological Processes (BPs) “platelet degranulation”, suggests that soluble mediators released after LPS treatment could bring platelets into the picture. A crucial factor for endothelial growth and repair is vascular endothelial growth factor (VEGF). It is known that platelets can participate in the restoration of tissue (i.e. endothelial) damage interacting with iPSC-MSC through the release of different factors [37]. Degranulation of platelets by factors released by treated iPSC-MSC may be of particular interest for endothelial repair, considering that platelets produce VEGF in many physiological situations, such as inflammation [38]. Although in our experimental model we did not include these cells, future research will be done in this way. It should be noted that levels of VEGF in ELISA assays from CM of treated iPSC-MSC were undetectable, even though these cells are known to produce and release this endothelial growth factor [39]. The release of VEGF through the exosome pathway, could be another mechanism to explain the lack of detection of this growth factor in our analyses [40], as our protocol for sample preparation for proteomic analysis did not break these vesicles [19].

In conclusion, in this work we showed that LPS generates an inflammatory program in iPSC-MSC that induces migration and adhesion to proteins present in ECM and these results was in concordance with the secretion of different proteins observed in ELISAs and proteomic assays. Stx alone did not induce inflammatory responses, even though iPSC-MSC expresses Gb_3_, but when combined with LPS, it decreased the capacity of endothelial cells to resolve a wound. The results observed in this work helps understand the role of iPSC-MSC in tissue regeneration, indicating that the immune context generated from these cells in response to a particular bacterial toxin should be taken into account.

## Data Availability

All LC-MS/MS raw data will be publicly available at Mass Spectrometry Interactive Virtual Environment (MassIVE) upon publication of the manuscript.

## Acknowledgments

The authors would like to thank LIAN (FLENI-CONICET) for allowing us to use laboratory facilities to perform many of the experiments from this work.

## Supplemental information

*Supp. Figure S1*. **LPS-dependent ICAM-1 expression in endothelial cells.** Mean fluorescence intensity (MFI) of ICAM-1 is shown for endothelial cells (HMEC-1) treated or not with 100 ng/ml LPS for 2 h. Results were expressed as mean ± S.E.M. n = 6 per group; **P<0.01.

*Supp. Figure S2*. **Biological processes related to proteins identified in CM of iPSC-MSC reflect the migration, adhesion to substrate and immune response induced by toxins.** (A) Heatmap shows normalized levels of proteins from collapsed technical replicates identified by proteomic analysis in the CM of untreated (Control, n=3) iPSC-MSC or treated with LPS (n=3), Stx (n=3) and LPS+Stx (n=3). Dendrograms represent the unsupervised euclidean (method) clustering of peptides (left). Names of identified proteins are shown to the right of the plot. (B) Overrepresented biological processes related to clustered proteins from (A) ordered by gene ratio (percentage of identified proteins in an ontology term). Size of spheres denotes the number of proteins contained in each process and color features the statistical significance of the algorithm represented by the adjusted p-value (Benjamini-Hochberg).

