## Supplementary figures and images for "Effects of bacterial lipopolysaccharide and Shiga Toxin on induced Pluripotent Stem Cell-derived Mesenchymal Stem Cells"

### Supplementary Figure 1

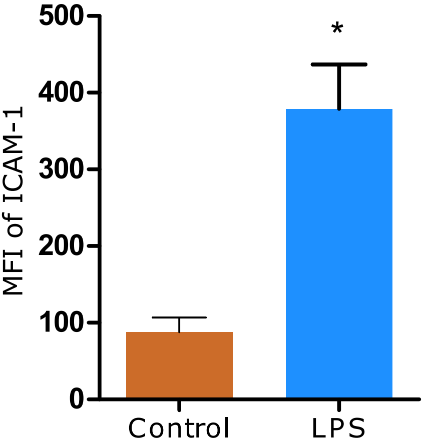

### Supplementary Figure 2

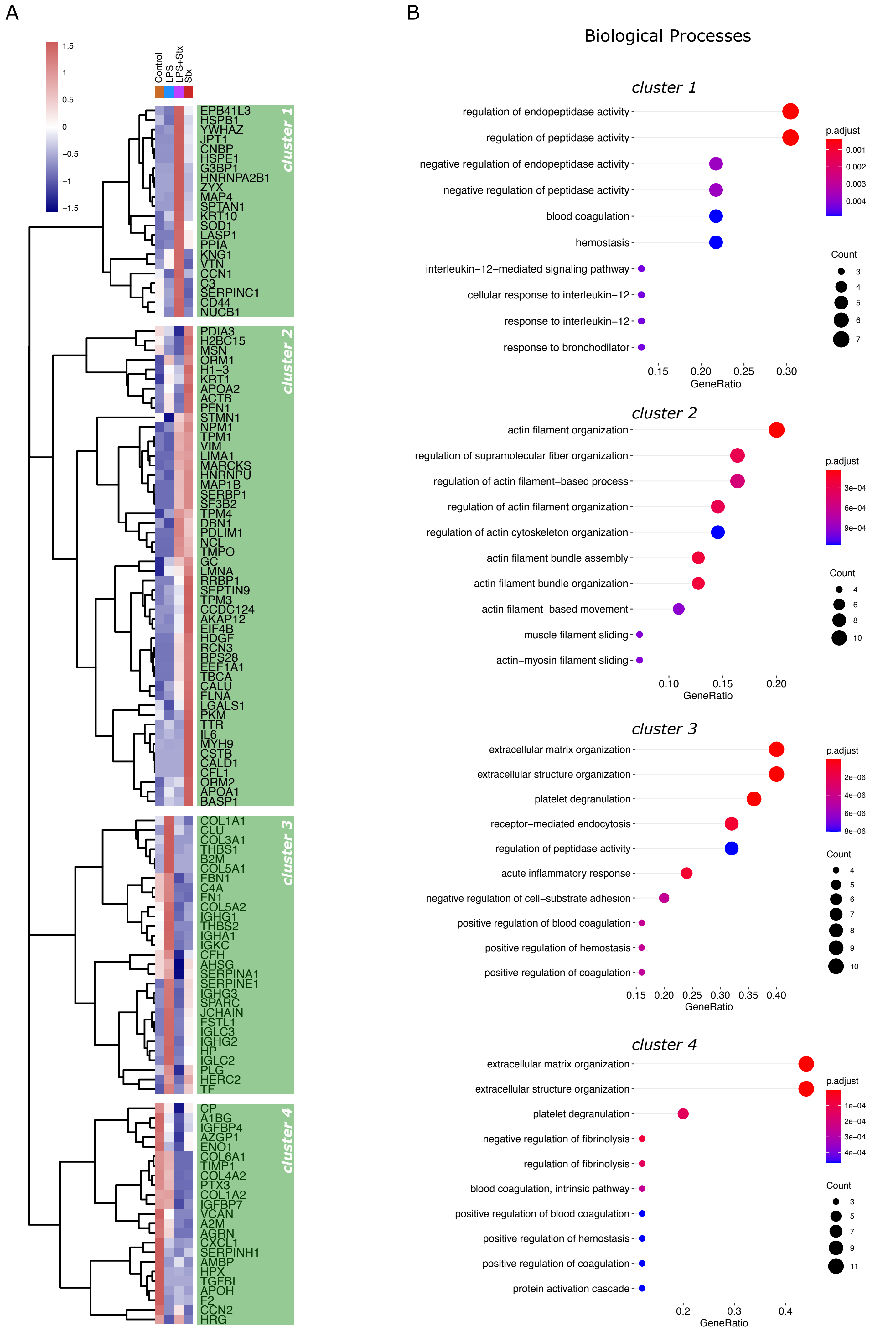
